# A comprehensive multicenter comparison of whole genome sequencing pipelines using a uniform tumor-normal sample pair

**DOI:** 10.1101/013177

**Authors:** Ivo Buchhalter, Barbara Hutter, Tyler S. Alioto, Timothy A. Beck, Paul C. Boutros, Benedikt Brors, Adam P. Butler, Sasithorn Chotewutmontri, Robert E. Denroche, Sophia Derdak, Nicolle Diessl, Lars Feuerbach, Akihiro Fujimoto, Susanne Gröbner, Marta Gut, Nicholas J. Harding, Michael Heinold, Lawrence E. Heisler, Jonathan Hinton, Natalie Jäger, David Jones, Rolf Kabbe, Andrey Korshunov, John D. McPherson, Andrew Menzies, Hidewaki Nakagawa, Christopher Previti, Keiran Raine, Paolo Ribeca, Sabine Schmidt, Rebecca Shepherd, Lucy Stebbings, Patrick S. Tarpey, Jon W. Teague, Laurie Tonon, David A. Wheeler, Liu Xi, Takafumi N. Yamaguchi, Anne-Sophie Sertier, Stefan M. Pfister, Peter J. Campbell, Matthias Schlesner, Peter Lichter, Roland Eils, Ivo G. Gut, David T. W. Jones, on behalf of the ICGC Verification and Validation Working Group

## Abstract

As next-generation sequencing becomes a clinical tool, a full understanding of the variables affecting sequencing analysis output is required. Through the International Cancer Genome Consortium (ICGC), we compared sequencing pipelines at five independent centers (CNAG, DKFZ, OICR, RIKEN and WTSI) using a single tumor-blood DNA pair. Analyses by each center and with one standardized algorithm revealed significant discrepancies. Although most pipelines performed well for coding mutations, library preparation methods and sequencing coverage metrics clearly influenced downstream results. PCR-free methods showed reduced GC-bias and more even coverage. Increasing sequencing depth to ∼100x (two- to three-fold higher than current standards) showed a benefit, as long as the tumor:control coverage ratio remained balanced. To become part of routine clinical care, high-throughput sequencing must be globally compatible and comparable. This benchmarking exercise has highlighted several fundamental parameters to consider in this regard, which will allow for better optimization and planning of both basic and translational studies.

## Introduction

‘Next-generation’ sequencing technologies have been available for almost a decade^1,2^, and in the past 5 years or so, their use in an ever increasing range of fields has resulted in an unprecedented revolution in genomic profiling (reviewed in^3^, for example). With the latest generation of sequencing machines promising to dramatically increase throughput and reduce costs, it is now inevitable that this technology will become a routine part of clinical care for a wide variety of diseases within the relatively near future. It is therefore critical to understand every stage of the analytical process in detail, from initial library generation (evenness of coverage, GC bias, introduction of artifacts, *etc*.) right through to annotation of the final variant lists. The question of ‘how much is enough’ sequence coverage to give sufficient power for genome-wide mutation detection has also not yet been conclusively addressed. The International Cancer Genome Consortium (ICGC^4^) has been one of the leading generators of cancer whole-genome sequencing (WGS) data in recent years. Under the auspices of this consortium, the Verification and Validation Working Group was established to investigate the factors that need to be considered in order to generate high-quality and high-confidence variant calls from WGS data. One method through which this has been evaluated is the establishment of two parallel benchmarking exercises to look at how differences in variant calling pipelines (Benchmark 1, BM1) and/or complete sequencing pipelines (Benchmark 2, BM2) can influence downstream results. We describe here BM2, where a single tumor-blood DNA pair was sequenced at multiple sites (the National Center for Genome Analysis (CNAG), Barcelona, Spain; the German Cancer Research Center (DKFZ), Heidelberg, Germany; the RIKEN institute, Tokyo, Japan; the Ontario Institute for Cancer Research (OICR), Toronto, Canada and the Wellcome Trust Sanger Institute, Hinxton, UK). The results were subsequently compared using both local and centralized analysis. The tumor chosen for this analysis was a medulloblastoma (a malignant pediatric brain tumor arising in the cerebellum^5,6^) from the ICGC PedBrain Tumor project. This tumor type typically shows a very high tumor purity (usually >95%), but also often carries ploidy changes and other copy number alterations, thereby allowing for analysis of mutation detection performance at different allele frequencies^7^. Merging data from the different contributing centers and analyzing the combined dataset resulted in an extremely high WGS coverage of >300x for the tumor and >250x for the germline control. This allowed us to investigate variant-calling parameters at very low allele frequencies, as well as the impact of imbalanced tumor *vs.* control coverage levels and of total sequencing coverage on mutation detection performance. These ‘real-life’, high-coverage tumor data complement a recent TCGA-ICGC benchmarking exercise on simulated data as part of the DREAM somatic mutation calling challenge^8^.

The results outlined below highlight a number of variables, some more surprising than others, as having an impact on the ultimate output of a sequencing experiment (*i.e.* the variant call list). Library preparation played a major role in output variability, with PCR-free libraries giving more even coverage and also higher coverage in regions of interest such as exons. Increasing coverage led to better mutation calling performance (with a notable proportion of calls missing at 30-40x), but only up to a saturation point at about 100x, and only when keeping tumor and control coverage roughly even. These findings raise important considerations for factors to take into account when planning future sequencing studies.

## Results

### Influence of library preparation on sequencing metrics

Several different protocols were used for generating sequencing libraries at the contributing centers, which varied in their reagent supplier, methods for selecting the fragment size of library inserts, and use of amplifying PCR steps (**Table 1**, **Online Methods** and **Supplementary Table 1**). Interestingly, these differences resulted in marked variation in the evenness of coverage genome-wide as well as in key regions of interest such as exons. Chromosome 22, being a small chromosome without copy number aberrations in this tumor (**Supplementary Figure 1**), was chosen to further assess base-wise coverage. The standard deviation of coverage ranged from 8.58 in the most evenly covered library to 38.49 in the most unevenly covered, which could have a significant impact on the ability to call copy number variations (**Supplementary Table 2**). Differences in coverage between tumor and control also influence the ability to call simple somatic mutations (SSMs) and small insertions/deletions (somatic indel mutations, SIMs), so we additionally calculated the standard deviation of the absolute pairwise coverage difference (tumor vs. control). The values ranged from 5.14 in a good library to 11.37 in a strongly biased library (**Supplementary Table 2**). PCR-free libraries were found to give the most even coverage, with very little effect of GC content on coverage levels, although several protocols containing an amplification step performed almost as well (**Figure 1a**). Two methods showed a marked variation in coverage, with a dramatic and unexpected increase in the number of sequencing reads mapping to regions of high GC content. This also resulted in much ‘noisier’ copy number profiles derived from these libraries, likely reducing the resolution at which structural variants could be reliably called (**Supplementary Figure 1**). One possible explanation for this maybe DNA-binding beads used during the clean-up process, which could feasibly bind more strongly to GC-rich sequences at a given fragment size under certain concentration and/or temperature conditions.

**Figure 1:**
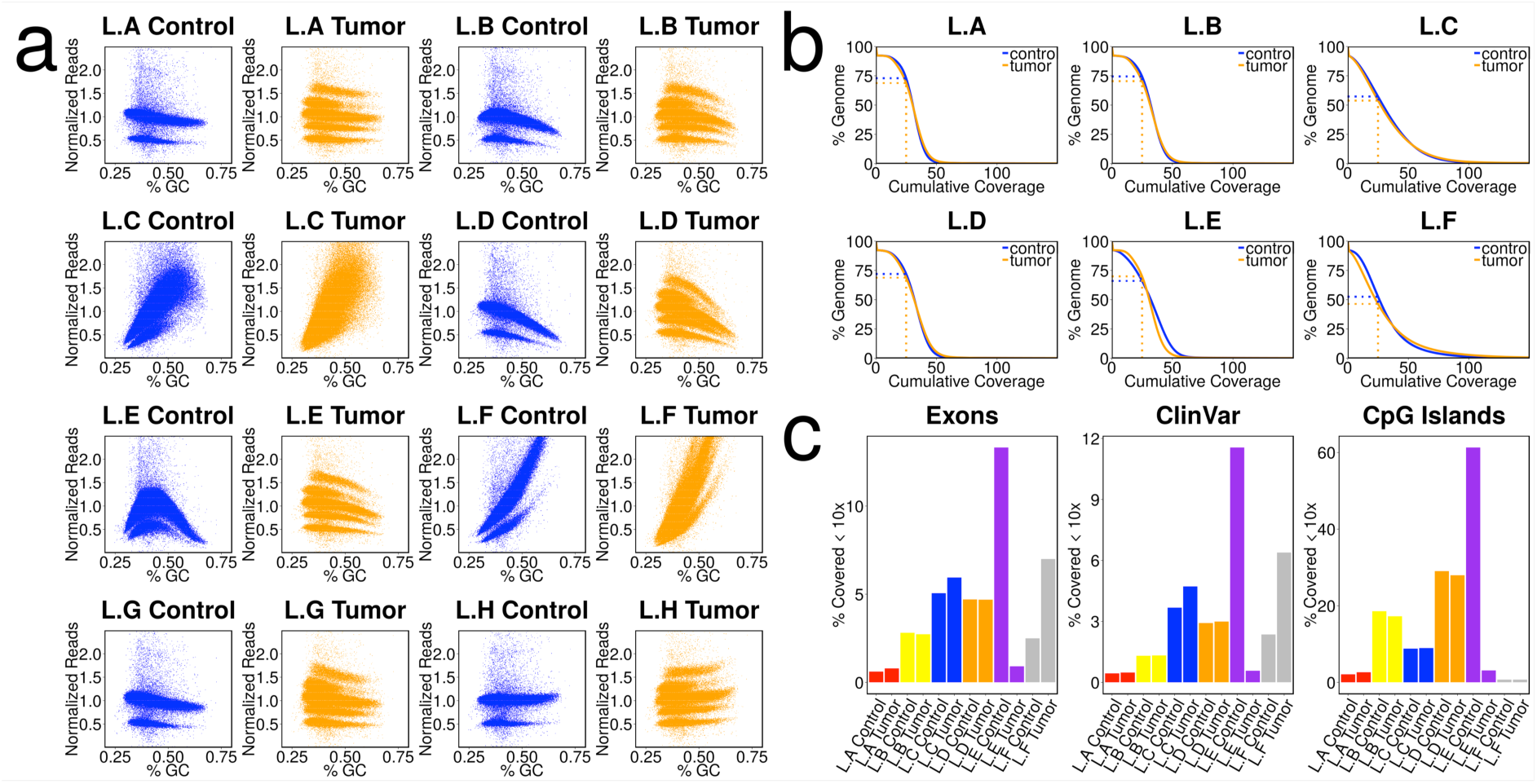
Differences between the different sample libraries. a) GC bias of the different libraries. The genome was segmented into 10 kb windows. For each window the GC content was calculated and the coverage for the respective library was added. For a better comparability the coverage was normalized by division with the mean. b) Cumulative coverage displayed for different libraries. Displayed are all libraries sequenced to at least 28x. To make the values comparable we downsampled all samples to a coverage of 28x (the lowest coverage of the initially sequenced libraries). The plot shows the percentage of the genome (y-axis) covered with a given minimum coverage (x-axis). c) Percentage of certain regions of interest covered with less than 10x.

**Table 1.**

| Library | Starting DNA (μg) | Fragment Size (bp) | Size selection | Library Protocol | PCR cycles | Sequencing Machine | Chemistry (Illumina) |
| --- | --- | --- | --- | --- | --- | --- | --- |
| L.A | 1 | ~ 400 | E-gel (Invitrogen, 2% Agarose) | TrueSeq DNA | 10 | HiSeq 2000 | V3 |
| L.B | 4 | ~ 400 | 2% Agarose Gel | KapaBio | no | HiSeq 2000 | V3 |
| L.C | 2.5 | ~ 500 | 2% Agarose Gel ~650 bp | NEBNext | 12 Cycles Post Ligation Clean Up | Both HiSeq 2500/2000 | v1 (RR)/v3 (HT) |
| L.D | 1 | ~ 550 | Agarose Gel 500-600bp cut | TrueSeq DNA | 10 | HiSeq 2000 | V3 |
| L.E | 2.8 | ~ 620 | 1.5% Agarose Gel (Pippin) | NEBNext | No | HiSeq 2000 | V3 |
| L.F | 1 | ~ 400 | AMPureXP Beads | NEBDNA | 10 | HiSeq 2000 | V3 |
| L.G | 1 | ~ 350 | AMPureXP Beads | TrueSeq DNA PCR-Free | 0 | HiSeq 2000 | V3 |
| L.H | 0.5 | ~ 175 | AMPureXP Beads | SureSelect WGS | 10 | HiSeq 2500 | V3 |

A better evenness of read distribution also means that a greater proportion of the genome is covered to a reasonable degree at a given average coverage level. It is preferable, for example, that the entire genome be covered at 30x rather than having half covered at 15x and half at 45x, even though both require the same total read number. To give a fair comparison for this measure, each dataset was down-sampled to an average 30x tumor and control coverage (the minimum coverage in any individual tumor or control set). In the best performing library, 74% of the genome was covered at or above 25x, while the worst performing library had only 46% of regions covered at this level (**Figure 1b**). In general the coverage distribution was more even and the percentage of well-covered regions was higher in the control libraries compared to the tumor libraries, reflecting the different copy number states of the tumor. An unusual pattern of GC content distribution in control library E, however, meant that this was slightly worse than its tumor counterpart.

The percentage of exonic regions covered at ≤10x (*i.e.* likely insufficient to accurately call mutations) also varied, with a range from less than 1% ‘missing’ in the best performing libraries to more than 10% in the worst (**Figure 1c**), demonstrating that sequencing library preparation performance can have a significant impact on the ability to identify variants in downstream analyses. Performance in other regions of interest, such as enhancers and UTRs, was similarly variable (**Figure 1c** & **Supplementary Figure 2**) While combining all libraries to give a coverage of close to 300x reduced the ‘missing’ exon fraction to just 0.1%, some regions of the genome (including part or all of ∼80 genes) were still only covered at <=10x (**Supplementary Table 3;** none of these genes are listed in the Cancer Gene Census^9^). The vast majority of these regions (>98%) were in non-uniquely mappable areas such as telomeric or centromeric repeats. These will likely never be covered using routine short-read methods, regardless of the total read count (*e.g.* in stretches of long, highly homologous repeats). Library A also contained some longer 2x 250bp MiSeq reads as opposed to standard HiSeq 2x 101bp, but the overall contribution of these (below 2x) was too low to assess whether they may help in covering some of the missed regions.

We also examined the performance of each dataset in regions of biased nucleotide composition that were previously reported to be challenging to sequence across different platforms ^10^. There was a marked variation in coverage in these regions, in keeping with the notable GC-bias observed in some libraries (**Supplementary Table 4**). The best overall performance in terms of evenness of coverage was seen with the PCR-free library, and this also outperformed the methods previously reported in the study of Ross *et al*. ^10^. Of note, some regions showed a significant discrepancy in coverage between tumor and normal in certain regions, which would likely compromise variant calling in these loci.

### Comparison of variant calling on the different libraries

The first comparison of variant calls that we performed was using each individual center’s own mutation calling algorithm on their sequencing output, which resulted in a surprising amount of variation. The precise differences in the variant calling algorithms are explored in more detail in the related ICGC Benchmark 1 exercise (Alioto *et al*., described elsewhere in this issue). Whilst there was a core set of mutations called by all 5 centers, this was the case for less than 20% of the total number of called variants (**Figure 2a**). Allele frequency plots indicated that these consensus calls showed clear peaks at ∼50% (heterozygous mutations occurring while the tumor was diploid) and ∼25% (mutations occurring in 1 of 4 alleles after tetraploidization of the genome), while those made by less than 4 centers were shifted towards a lower allele frequency. This may indicate either increased variability in sensitivity of the pipelines as allele frequency decreases, and/or some mutations at such a low frequency that there were no variant reads in certain datasets (**Figure 2a**). The mutation contexts of the variants were reasonably similar across centers, with the majority being C > T transitions in a GpCpG or ApCpG context, although some variability can clearly be seen across the 5 sets (**Figure 2b**). Roughly one third of the mutations were unique to only one center, with the remainder variably called by 2-4 groups. One of the most notable differences was the low total number of calls made by center C, resulting in a large proportion of calls called by the other 4 centers but not this one. Based on the outcome of the ICGC benchmark analyses, however, this center has now modified its analysis pipeline to slightly relax some over-stringent filtering steps, resulting in a much greater overlap with the other calls (not shown). When looking further at mutational signatures as defined by Alexandrov and colleagues^11^ rather than simple base change contexts, variation can also be seen per center in the number and type of mutational processes identified (**Figure 2c**).

**Figure 2:**
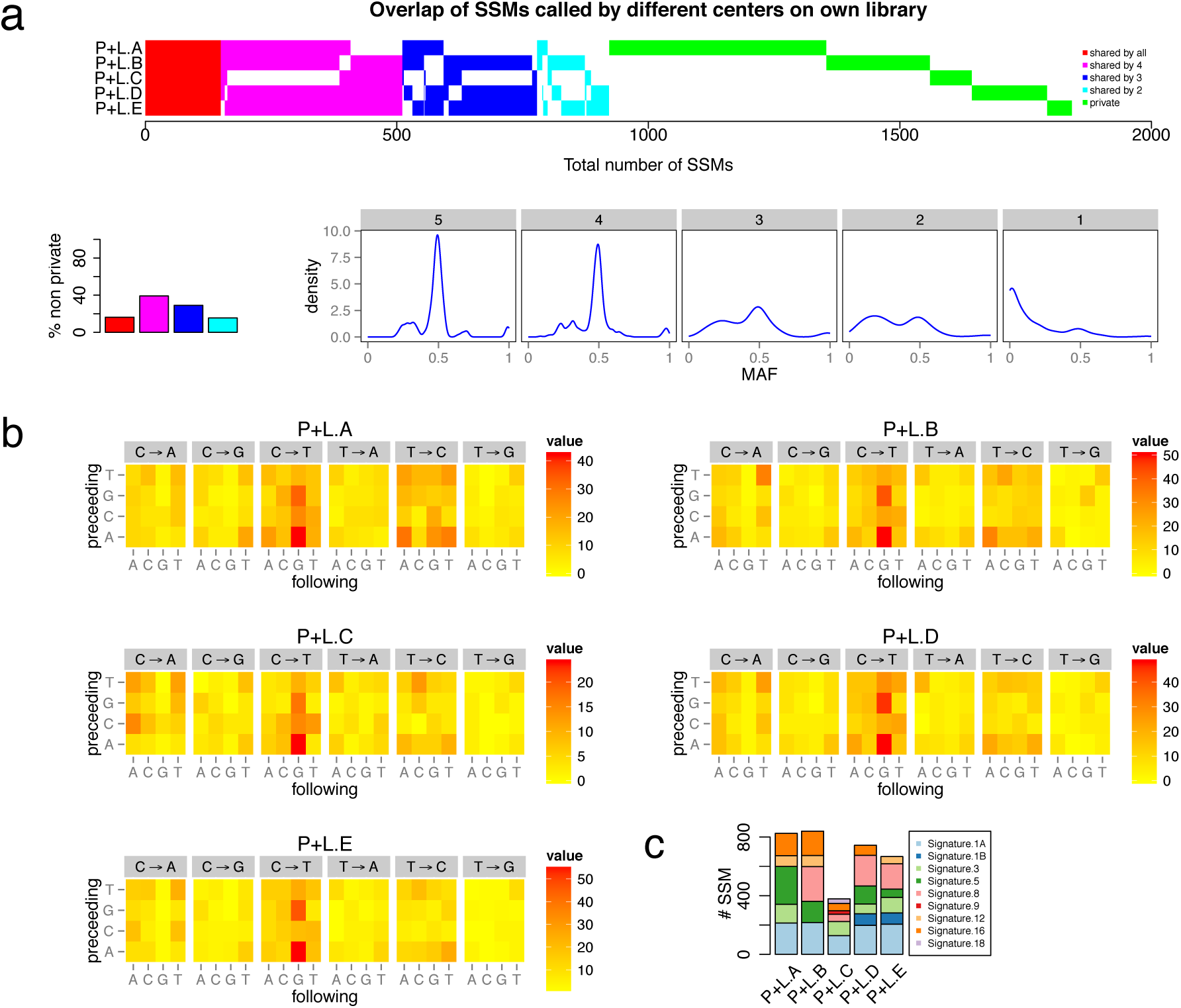
Comparison of ability to discover SSMs with different pipelines. a) Overlap of SSMs called by each center on its own library. All SSMs detected by at least one center are shown on the x-axis. The SSMs were sorted and colored by recurrence. SSMs were considered to be identical when both, the exact position and the base substitution were the same. The bar plot shows the percentage of all non-unique SSMs for the given levels of concordance. Shown on the bottom are the density plots of the variant allele frequencies for each level of concordance. b) Sequence context of SSMs detected by each center on its own library. For each single base substitution, the sequence context (plus/minus one base) was determined. The 128 possible combinations are shown in a heat map. c) Mutational signatures for SSMs as defined by Alexandrov and colleagues^11^. The calls from each center were used to fit into the predefined signatures. Only signatures composing at least 5% of the total SSMs are shown.

In terms of coding alterations, there was a greater degree of overlap, but certainly not 100% concordance. A curated ‘Gold’ set of calls was generated from the 300x WGS dataset by comparison of multiple independent call sets followed by manual inspection (Alioto *et al*., described elsewhere in this issue and available from the ICGC Data Portal, https://dcc.icgc.org/). Four non-synonymous, one splice-site (*SET*) and one stop gain (*ANGPT1*) SSMs were identified from this Gold set, which were also present at more than 10% allele frequency in each individual dataset. Of these, one center called all 6, two centers called 5, and one 4. One outlier called only two originally, which was found to be a result of minor contamination of the control sample with tumor DNA. A second library preparation resolved this issue, and all 6 SSMs were subsequently called (**Supplementary Figure 3**). One analysis pipeline also indicated a potential SSM in *ZMYM3* that was not detected in the other sets. Further inspection revealed that this alteration is probably a complex SIM rather than a single point change (discussed below).

Interestingly, the variation of calls between these five centers was higher for this exercise than for the Benchmark 1 exercises (Alioto *et al*., described elsewhere in this issue). In particular, each center calling on their own library produced a higher variation than for the same centers calling on the same tumor-normal pair, but on data from only one center (Benchmark 1.2), clearly indicating that library variations contribute to the observed heterogeneity of mutation calls. When excluding unique calls, fewer than 60% of SSM calls were shared between 4 or more centers and fewer than 20% were called by all 5 when analyzing different libraries. When using only one library, however, more than 60% of SSM calls were shared across all centers (**Supplementary Figure 4**).

Although the previous comparison already provided some evidence of a role for pre-analysis sequencing pipelines in generating differences between datasets, we wanted to further assess this by removing any variation in the analysis pipeline itself. We therefore re-aligned and re-called mutations on each dataset using one standardized pipeline (the DKFZ pipeline was chosen for logistical reasons). This resulted in a notably better consensus of mutations called by more than one center (>80% called by at least 4 out of 5 centers, **Figure 3a**, versus <60% with different pipelines, **Figure 2a**), but an unexpected increase in the number of private mutations, particularly for one center (**Figure 3a**). A shift towards lower allele frequencies was again seen in the mutations not called in all centers (**Figure 3a**). Analysis of mutation contexts indicated that the vast majority of these excess mutations were T>G transversions with low allele frequency, which were not observed at high frequency in the other datasets. Simply filtering out mutations with low allele frequency arising in this context resulted in an improvement in the overlap of mutation calls, but many more exclusively called (‘private’) alterations remained compared with the center’s own calls on their data (filtered against other reference samples, **Figure 2a** and **Figure 3b**). Closer investigation revealed that the cause for this artifact was a center-specific method for adjusting base quality q-scores, whereby a calibrating PhiX library was spiked into each sequencing lane. Unfortunately, this actually led to an increase in the specific artifact detected in this comparison, and the center has subsequently reverted to default q-score metrics. The fact that the same phenomenon was not seen in the center’s own calls on their data (**Figure 2**) is because it had already been identified, and a customized filter applied to account for it (removal of such changes also observed in a panel of 48 sequenced normal samples). This emphasizes that care must be taken when re-analyzing publicly available genome data from external centers, particularly when details on library preparation and customized ‘blacklists’ are not known. This effect also had an impact on the mutational signatures identified, with a different distribution of processes observed in this mutation set than for each center calling their own variants (**Figure 3d**), further suggesting that both library preparation and calling algorithms can strongly affect the ability to accurately detect such signature.

**Figure 3:**
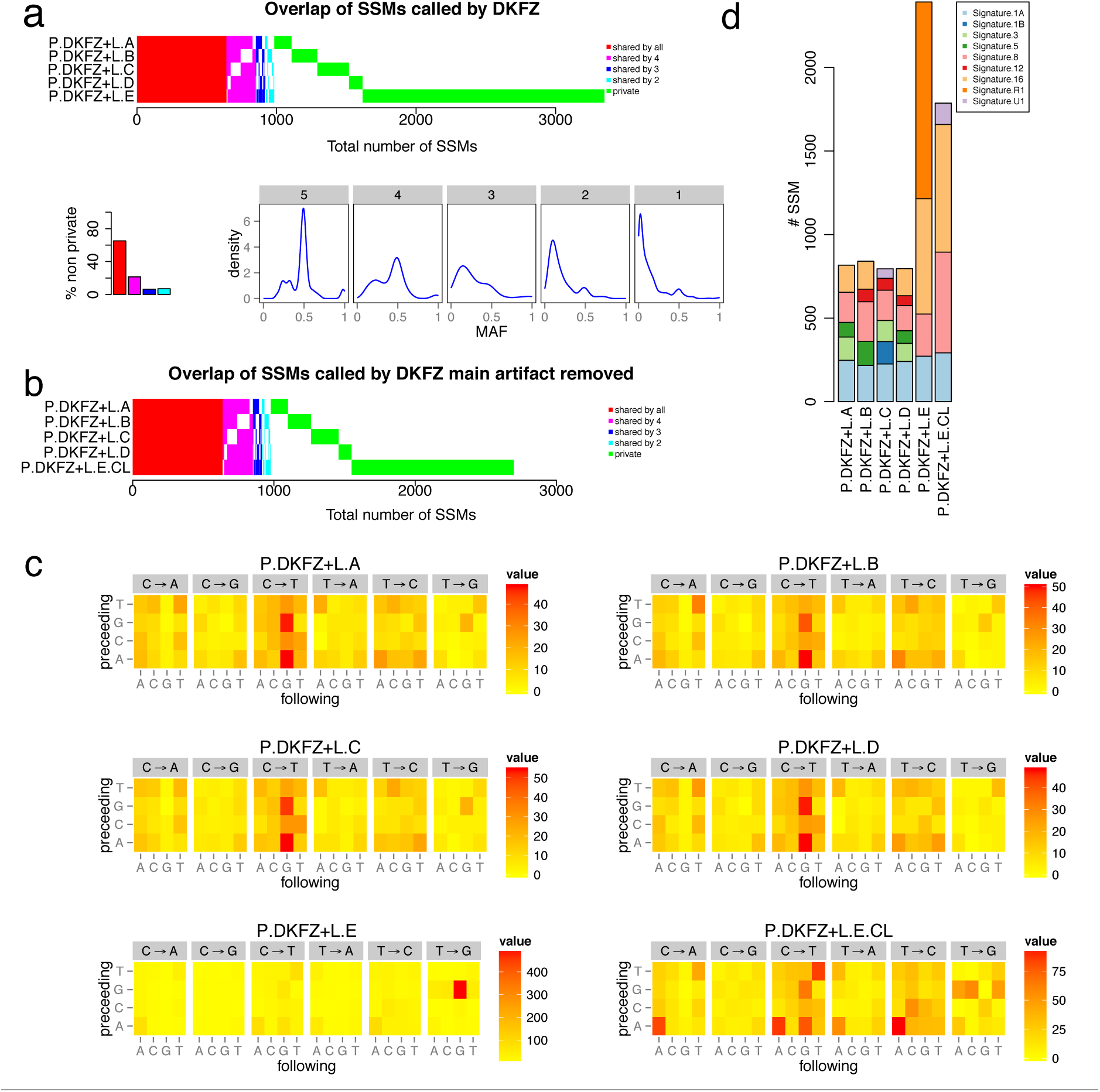
Influence of the library and sequencing on SSM calls with one pipeline. a) Overlap of SSMs called using one pipeline (DKFZ) on all different data sets. Percentage of concordance of non-unique SSMs are shown in the bar plot. The bottom shows density plots of variant allele frequencies for each concordance level. b) Overlap of SSMs called after removal of the most prominent artifact (GpTpG to GpGpG) from the library L.E. c) Sequence context of SSM calls derived from the DKFZ pipeline on the different data sets. In order to have a better comparability, the most prominent artifact was removed from L.E (L.E.CL). d) Mutational signatures for SSMs as defined by Alexandrov and colleagues^11^. Calls made by DKFZ were fitted to the predefined signatures. Only signatures composing at least 5% of the total SSMs are shown.

### Effects of tumor/normal coverage levels on variant calling

Combining the sequencing data generated from each participating center gave us the excellent opportunity to investigate a tumor-normal pair with very deep coverage whole-genome sequencing information. After merging each of the individual pairs, the combined tumor coverage was 314x, and the control 272x. To remove already identified artifacts, we excluded the tumor library from center E and the slightly contaminated control library from center B. For comparison of mutation calling metrics at a range of coverage levels, the combined tumor and normal sets were randomly serially down-sampled to 250, 200, 150, 100, 50, 30 and 20x coverages and then analyzed using the standard DKFZ pipeline. The total number of mutations increased when going from 30x to 50x and further to 100x coverage, but no striking increase was seen above this level (by 100x, 95% of the maximum mutation number are detected, in contrast to only 77% at 30x; **Figure 4a, Supplementary Table 5**). Whilst the majority of mutations were called at the 30x level, there were some notable differences in the number and type of mutations detected as the coverage increased. In particular, the sensitivity for detecting mutations with lower mutant allele frequencies (*i.e.* subclonal alterations and/or events happening after polysomic changes but also major somatic mutations in samples with low tumor cell content) was much greater with higher coverages, as seen from density plots of mutations vs allele frequency (**Figure 4b**). This effect was even more striking when looking at mutation calls per chromosome, which clearly shows the difference between low and high coverages when looking for late-occurring mutations after whole chromosome copy number changes (**Supplementary Figure 5**).

**Figure 4:**
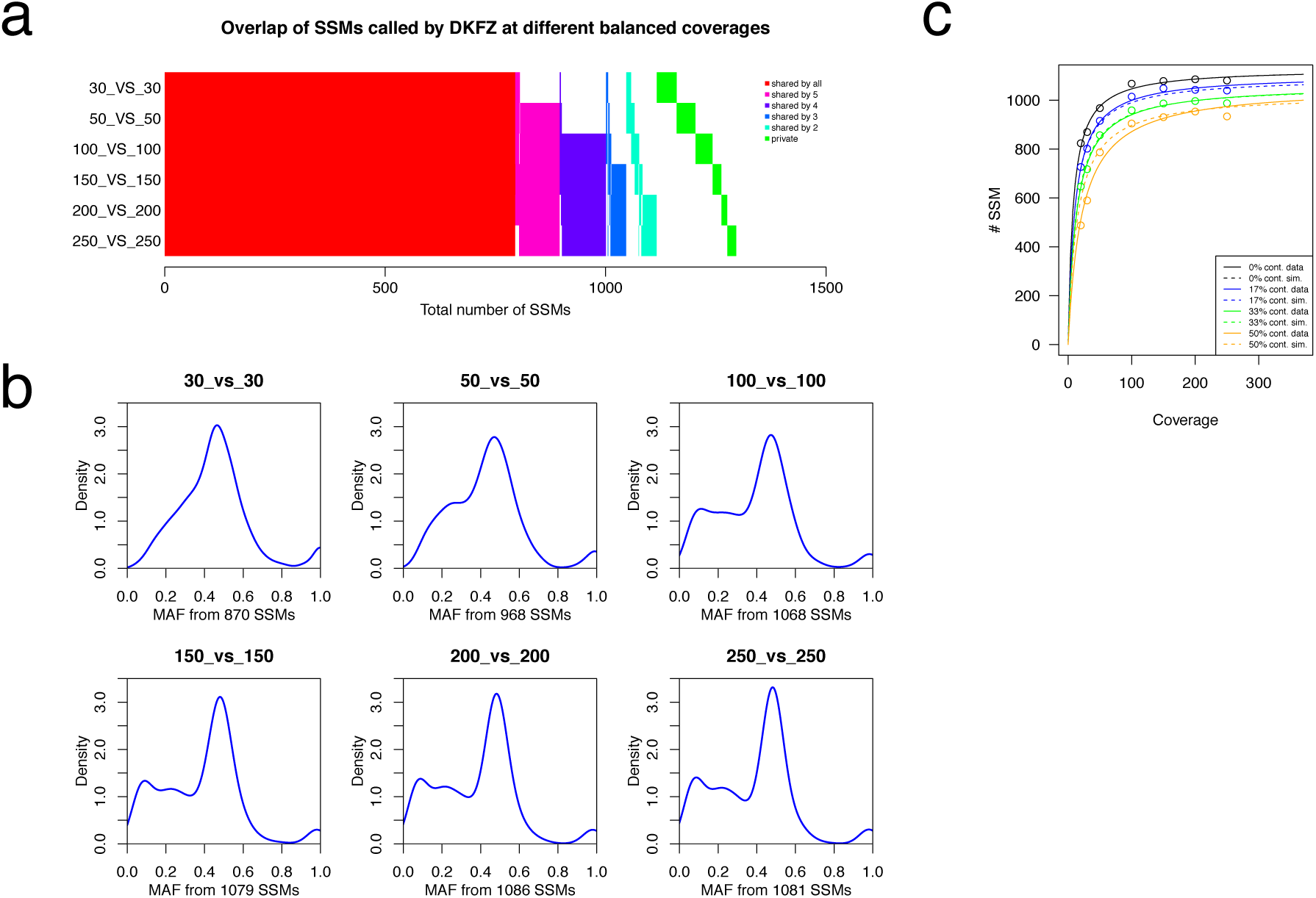
Effect of sequencing coverage (from downsampled combined files) on the ability to call SSMs. a) Overlap of SSMs called on different balanced coverages. b) Density plots of the variant allele frequencies for different balanced coverages of tumor and control (tumor_vs_control) and number of SSMs called in total (calls were done using the DKFZ calling pipeline).c) Plot of the number of SSMs (y-axis) found for a given coverage (x-axis). The different colors represent different levels of normal “contamination” in the tumor (0% black, 17% blue, 33% green and 50% orange). Solid lines represent the real data and dashed lines are simulated. Lines are fitted against the Michaelis-Menten model using the ‘drc’ package in R. Solid lines are fitted to the data points and dashed lines are simulated using a mixed inhibition model for enzyme kinetics.

Since medulloblastomas tend to show a very high tumor cell content (usually above 95%, and for this samples ∼98%, due to their nature as masses of small, round, tightly-packed tumor cells), the high coverage dataset also provided a good opportunity to model the dynamics of mutation calling with increasing coverage and with increasing proportions of ‘contaminating’ normal tissue (low tumor purity). We found that the mutation calls with increasing coverage were accurately modeled by a Michaelis-Menten equation, reaching ‘saturation’ (no or minimal additional mutations called as coverage increases) at around 100x (**Figure 4c**). For SIMs (indels) called using the DKFZ pipeline, a different picture was observed. SIM calling at present likely suffers at least much from low specificity as from low sensitivity, as indicated by the fact that increasing coverage actually reduces the number of called variants (i.e. the false positive rate decreases; **Supplementary Figure 6**). The impact of normal cells on SSM detection could be thought of as a ‘mixed-type inhibition’ of mutation detection sensitivity, which we examined by mixing increasing proportions of normal sequence reads (17%, 33% and 50%) into the tumor dataset and re-calling mutations. Each curve displayed the same plateau after ∼100x as the pure tumor sample, but the addition of any normal content meant that the maximum mutation count from the pure tumor could not be reached, even at 250x total coverage. At 100x, the detected proportion of mutation calls from the pure sample were 95%, 90% and 85% respectively for 17%, 33% and 50% ‘contamination’ (**Figure 4c**). At lower coverages, however, the normal cell content had a proportionally larger impact. At 30x, only 92%, 83% or 68% of the calls from the 30x pure sample were called when adding 17%, 33% or 50% normal reads, respectively (**Supplementary Table 5**).

We next investigated the effect of tumor:normal coverage ratios on variant calling, to assess whether increasing coverage of the tumor alone is sufficient to increase mutation detection sensitivity. The 250x tumor genome was therefore compared with a down-sampled control at 200, 150, 100, 50 and 30x coverages. Down to the 150x level, few differences are seen in the mutations called when compared with the 250x/250x standard (**Figure 5a,b**). At lower control coverage levels, however, a notable increase is observed in the overall number of mutations reported, due to a sharp rise in those called with a low allele fraction. Since these mutations are not called in the 250x vs 250x set, it is almost certain that they are sequencing artefacts arising in a very small proportion of calls, which appear to be somatic when the control coverage is insufficient to show the same phenomenon. When looking further into the context of these new calls, it is clear that they are dominated by one specific base change (T>G) arising in a particular sequence context (GpTpG, **Figure 5c**). Indeed, performing a motif analysis on the wider context of these changes revealed that the majority arise at a thymine base within a homopolymer run of guanines (**Figure 5d**). Keeping the ratio of tumor:normal coverage closer to 1 therefore appears to play a role in maintaining the accuracy of mutation calling with standard pipelines, since any systematic artifacts are then balanced out in both the tumor and control datasets. While it may be possible to apply additional filters to account for the new false positives seen in unbalanced comparisons, this would potentially come at the cost of a reduced sensitivity for detecting true mutations with low allele frequencies (*i.e.* tumor subpopulations), which are of particular interest when increasing sequencing coverage depth.

**Figure 5:**
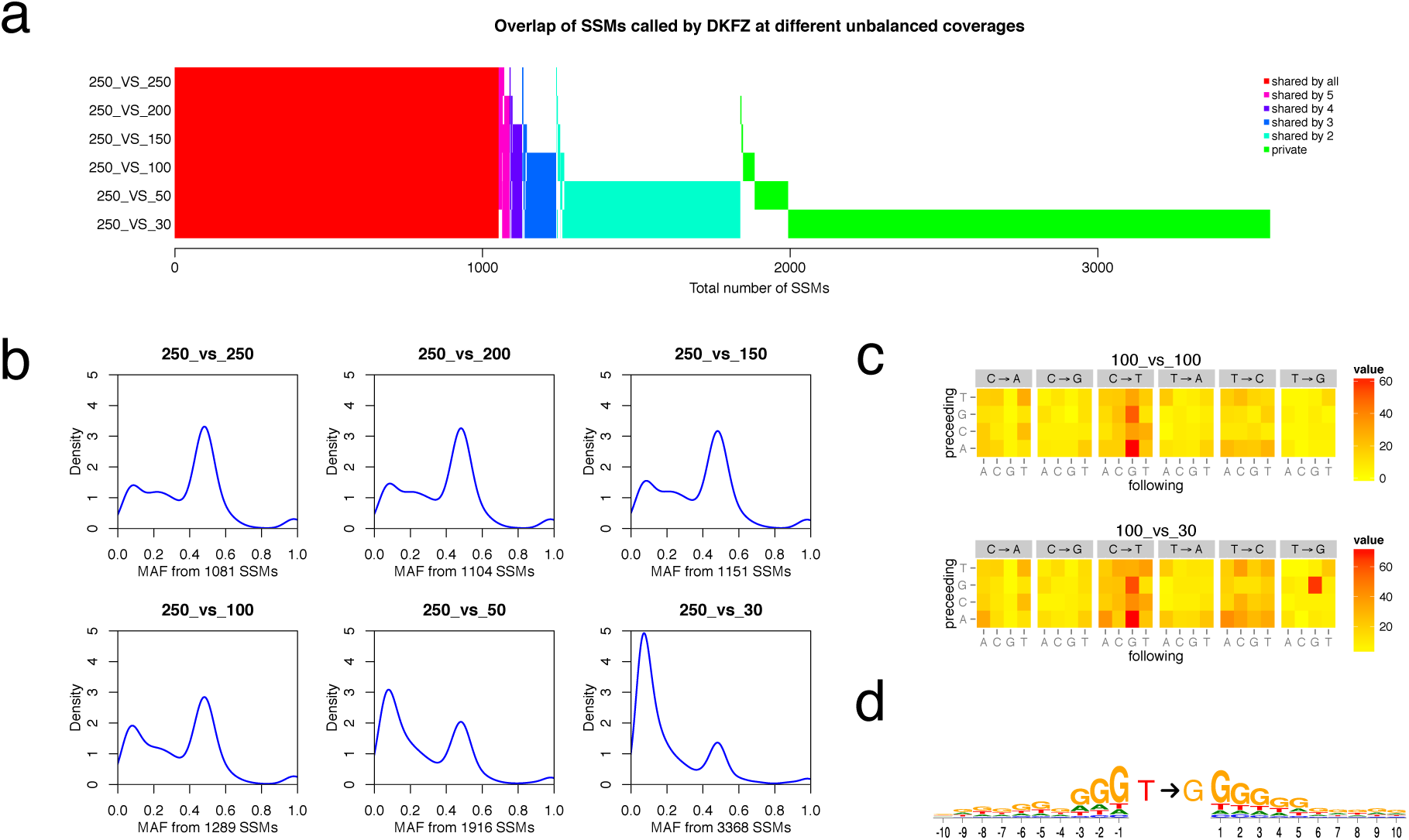
Effect of unbalanced coverage between tumor and control on SSM calling. a) Overlap of SSMs called on different unbalanced coverages. b) Density plots of the variant allele frequencies for different control coverages and a fixed tumor coverage and number of SSMs called in total (calls were done using the DKFZ calling pipeline). c) Sequence context of SSM calls derived for two different coverage combinations (100x tumor vs. 100x control and 100x tumor vs. 30x control). d) Logo plots showing the window of ten bases upstream and downstream from the presumed T to G transversion artifact.

## Discussion

This benchmarking exercise has highlighted the importance of carefully considering all stages of the laboratory and analysis pipelines required to run a sequencing experiment, in order to generate consistent and high-quality genomic data. Somewhat reassuringly for present applications, most of the coding mutations present at >10% allele frequency (which could reasonably be expected to be detected at ∼30x coverage) were accurately detected, especially after a second subsequent to optimization and harmonization of variant calling algorithms. The overlap was still not 100%, however. The problem of ‘missing’ SSMs was recently highlighted in colorectal and endometrial tumors, whereby frequent mutations were identified in *RNF43* that had not been detected in a previous analysis due to the sequence context^12^. Whilst likely not truly an SSM, and therefore not directly related to the current analysis, the almost coincidental finding of a *ZMYM3* alteration in the MB sample further highlights that some classes of alteration are poorly detected by current pipelines. This is particularly important when considering the potentially driving role of this gene in medulloblastoma^13,14^.

Discrepancies outside of the coding regions were more substantial, which may be of significance as the role of the non-coding genome to human disease becomes clearer^15,16^. Library preparation methods clearly had a significant impact on downstream data output, even when using one standardized variant calling pipeline. PCR-free libraries gave a more even coverage and also covered a higher proportion of functional regions such as exons than those requiring an amplification step. Each method was tested only once, however, so the reproducibility of variations and precise contribution of different parameters in the sequencing process could not be directly tested. The reporting of detailed protocols and QC metrics is therefore a necessity for clinical or other regulated applications^17^. Comparison of different coverage levels indicated that care should be taken not to increase artefact rates when increasing tumor but not control sequencing. The rate of new mutations identified with increasing coverage up to as much as 100x suggested that the 30-40x which is still often taken as standard^18^ may not be sufficient to capture the full spectrum of (especially non-coding) changes, particularly when there is an additional contribution of uneven coverage, low tumor purity or subclonal heterogeneity (although our method of merging data from multiple centers may not completely reflect deep coverage data from one center). Indeed, even at 250x coverage, not all mutations from the pure tumor could be recaptured from the samples ‘contaminated’ with normal DNA. This effect will most probably be more or less noticeable with different variant calling algorithms that have been optimized to work on the specific tumor types or sequencing protocols that are most frequently encountered at a given center (with varying inherent tumor purities and other types of bias), but it is likely that 30-40x is insufficient to truly capture the full somatic mutational spectrum of a sample, regardless of other parameters.

Taken together, our results suggest that PCR-free library preparation protocols should be the method of choice in order to ensure evenness of coverage, and that a sequencing depth of close to 100x for both tumor and normal ought to be aimed for (particularly in situations where subclonal mutations or non-coding alterations are suspected to be playing a role). With platforms such as the Illumina HiSeq X now coming online in more centers, such an increase in coverage may be feasible without dramatically increasing costs. We would also recommend that variant calling pipelines should be benchmarked against publicly available datasets of validated mutations, including the Gold set of mutations derived from the data presented here.

In summary, this valuable resource can serve as a useful tool for the comparative assessment of sequencing pipelines, and gives important new insights into sequencing and analysis strategies as we move into the next big expansion phase of the high-throughput sequencing era.

## Acknowledgements

We thank the DKFZ Genomics and Proteomics Core Facility and the OICR Genome Technologies Platform for provision of sequencing services. This work was supported by: the PedBrain Tumor Project contributing to the International Cancer Genome Consortium, funded by German Cancer Aid (109252) and by the German Federal Ministry of Education and Research (BMBF, grants #01KU1201A, MedSys #0315416C and NGFNplus #01GS0883); the Ontario Institute for Cancer Research to PCB and JDM through funding provided by the Government of Ontario, Ministry of Research and Innovation; Genome Canada; the Canada Foundation for Innovation and Prostate Cancer Canada with funding from the Movember Foundation (PCB). PCB was also supported by a Terry Fox Research Institute New Investigator Award, a CIHR New Investigator Award and a Genome Canada Large-Scale Applied Project Contract. Financial support was provided by the consortium projects READNA under grant agreement FP7 Health-F4-2008-201418, ESGI under grant agreement 262055, GEUVADIS under grant agreement 261123 of the European Commission Framework Programme 7, ICGC-CLL through the Spanish Ministry of Science and Innovation (MICINN), the Instituto de Salud Carlos III (ISCIII) and the Generalitat de Catalunya. SD is supported by the Parc Científic de Barcelona through the Torres Quevedo subprogram (MICINN) under grant agreement PTQ-12-05391. The Synergie Lyon Cancer platform has received support from the French National Institute of Cancer (INCa). The ICGC RIKEN study was supported partially by RIKEN President’s Fund 2011, and the supercomputing resource for the RIKEN study was provided by the Human Genome Center, University of Tokyo.

## Author Information

Sequence data are available from the European Bioinformatcs Institute (EBI) European Genome-phenome archive (EGA) under accession number EGAS00001001107.

## Methods

### Patient material

An Institutional Review Board ethical vote (Medical Faculty of the University of Heidelberg) as well as informed consent was obtained according to ICGC guidelines (www.icgc.org).

### Library preparation and sequencing

The libraries were prepared at the different sequencing centers. Some samples are the result of a mixture of different libraries (as per the center’s standard protocols); others are comprised of one library only. An overview of the composition of the different samples and differences in the library preparation protocols is given in **Table 1** and **Supplementary Table 1**. All samples were sequenced using Illumina technology and chemistry. The majority of reads are of 2x100 bp length and are derived from HiSeq2000 or HiSeq2500 sequencers. Only library L.A additionally has a low number of 2x250 bp MiSeq reads included.

### Comparison of SSM calls

Each of the participating centers performed mutation calling using the respective in house pipelines (Alioto T et al., accompanying manuscript). The raw simple somatic mutation (SSM) calls were provided in the form of customized variant calling files (VCF). In order to provide a fair comparison, only single base point mutations were considered. A call was considered to be equal when both the position and the exact substitution reported were identical. The calls were then sorted according to the number of centers that made this particular call using a custom Perl script. The resulting file was plotted using a custom R-script (both available on request).

### Merging of the bam files to get the 300x files

To create the high coverage ∼300x bam files, the raw fastq files were aligned using bwa 0.6.2-r126-tpx aln -t 12 -q 20. Followed by bwa-0.6.2-tpx sampe -P -T -t 8 -a 1000 -r. The bam files for each center/library were merged and duplicates were marked using Picard tools MarkDuplicates Version 1.61. Finally, all merged per center bam files were merged using picard-1.95 MergeSamFiles and the header was adjusted using samtools-0.1.19 reheader. Since only reads from different libraries were merged at this step, duplicates were not marked. The coverage was calculated using an in-house tool, taking into account only non-N bases.

### Downsampling of the 300x files

The ∼300x bam files were serially down-sampled to different coverage levels (250x, 200x, 150x, 100x, 50x, 30x, 20x) using picard-1.95 DownsampleSam, and the coverage was determined after each step.

### Determination of library GC-bias

To determine the GC-bias of the libraries we first created 10kb windows over the whole genome using bedtools (v2.16.2) makewindows. Then the GC content for each window was calculated using bedtools (v2.16.2) nuc, windows containing more than 100 “N” bases were excluded (awk-3.1.6 ‘BEGIN{FS=“\t”}{if ($10 <= 100 && $11 <= 100) print $1“\t”$2“\t”$3“\t”$5}’). Finally the coverage for each of the remaining windows was calculated using bedtools (v2.16.2) multicov. Since the total coverage of the different libraries was not the same, the coverage was normalized by diving the coverage for each window by the mean coverage across all windows for each of the samples respectively. To visualize the GC-bias we then plotted the normalized coverage against the GC-content.

### Determination of percentage of bases covered with fewer than ten reads in special regions of interest

The regions of interest were defined as previously described^19^. To determine the percentage of bases covered with fewer than ten reads, we first determined the coverage over the whole genome in per base resolution using genomeCoverageBed (v2.16.2) -bga. The resulting coverage file was compressed using bgzip and an index was produced with tabix-0.2.5 -p bed. We then extracted the coverage for our regions of interest using tabix-0.2.5. From the resulting extracted coverage files we computed the number of bases covered by a certain number of reads using intersectBed and a custom perl script. This table was then used to determine the percentage of bases covered by <= 10 reads.

### Extracting mutation signatures

Mutational catalogues were generated based on the somatic mutations detected in the tumors. The 3’ and 5’ sequence context of all selected mutations was extracted, and the resulting trinucleotides were converted to the pyrimidine context of the substituted base. Considering 6 basic substitution types with surrounding sequence context, this results in a mutation type vector of length 96. The mutational catalogue was set up by counting the occurrence of each of these 96 mutation types per sample.

The proportions of the signatures published by Alexandrov *et al*.^11,20^ contributing to the mutational profile of each sample were estimated based on the probabilities of point mutations with their trinucleotide context in the signatures. The respective exposures were extracted sample-wise by quadratic programming. Exposures were plotted if they accounted for at least 5% of the SSMs in a sample.

